# Tracking genome evolution in single cell clones reveals the rates and features of copy number alterations generated by ongoing chromosomal instability in cancer

**DOI:** 10.1101/2023.09.27.559836

**Authors:** Molly A. Guscott, David Gómez-Peregrina, Alexander Malling Andersen, Tanya N. Soliman, Caterina Vidal Horrach, Bjorn Bakker, Diana Carolina Johanna Spierings, René Wardenaar, Floris Foijer, Cesar Serrano, Roland Schwarz, Sarah E. McClelland

## Abstract

Cancer genomes exhibit extensive chromosomal alterations caused by ongoing Chromosomal Instability (CIN). The ensuing cell-cell heterogeneity facilitates evolution and cancer cell plasticity that can drive therapy resistance, yet cancer CIN driver mechanisms remain essentially uncharacterised. This lack of knowledge presents an untapped opportunity to target vulnerabilities associated with ongoing CIN for therapy. Existing methods to investigate the cellular mechanisms responsible for CIN rely on laborious functional assays, or inference from genomic alteration patterns from sequencing data. Current bulk sequencing derived copy number alteration pattern signatures lack the cell-cell resolution that would reveal recent genomic alterations caused by CIN. Large-scale single cell sequencing of cancer cell populations is now emerging. However, it is not known whether the effects of selection still obscure the spectrum of genomic alterations caused by recent CIN. To address this, we employed a single-cell whole-genome sequencing (scWGS) clonal outgrowth technique, that allows us to track the real-time evolution of cancer genomes at the single-cell level. Single cancer cells surprisingly re-establish heterogeneity that matches their parental population within ∼22 generations. By comparing the features of copy number alterations at different evolutionary timepoints we reveal that some alteration types are likely under negative selection and are thus only apparent in the most recent cell divisions, and not in the parental population. In one cell line we identify a particular chromosome subject to recurrent chromosomal deletions, and validated that this chromosome was involved frequently in mis-segregation events during anaphase using fluorescence *In-Situ* hybridisation.

## Introduction

Chromosome Instability (CIN) is a complex and poorly understood hallmark of cancer, characterised by continuous gains and losses of whole or parts of chromosomes during cell division, leading to highly diverse tumour populations. We and others have previously demonstrated using functional assays in cancer cell lines that multiple mechanisms can drive CIN. For example, replication stress and deviant microtubule dynamics have been observed in colorectal and high grade serous ovarian cancers (Burrell *et al*., 2013; Tamura *et al*., 2020b; Ertych *et al*., 2014; Thompson and Compton, 2011; Bakhoum, Genovese and Compton, 2009). However, cell biological assays are limited in their resolution, meaning precise mechanisms of CIN remain elusive.

Chromosomal instability results from disregulation of essential genome stability pathways, leaving cancer cells vulnerable to synergistic therapies. For example, ATR inhibitor AZD6738 is effective against high replication stress cancer cells, while PARP inhibitors act upon tumours with homologous recombination deficiency (HRD) (Forment and O’Connor, 2018; Bryant *et al*., 2005; Farmer *et al*., 2005; Ubhi and Brown, 2019). Recent studies have unveiled characteristic genomic alteration patterns resulting from chromosomal instability processes (Shaikh *et al*., 2022b), offering an opportunity for tailored cancer treatments based on individual tumour genomic alterations. HRD genomic “scars” in BRCA-deficient tumours serve as a prime example, guiding the use of PARP inhibitors in clinical decisions.

Two significant challenges hinder the extension of this principle to new successful cancer treatment strategies. First, our understanding of chromosomal instability mechanisms in cancer remains incomplete and is currently limited to low-throughput functional analyses in cancer cell lines. Second, pinpointing the specific CIN mechanism at work in a patient’s tumour remains elusive. Recent studies have attempted to identify CIN drivers in cancer through computational “CIN signatures” based on DNA copy number alteration (CNA) patterns observed in bulk whole genome sequencing of tumours (Davies *et al*., 2017; Nik-Zainal *et al*., 2016; Macintyre *et al*., 2018; Ng *et al*., 2012; McBride *et al*., 2012). These represent a parallel to the single base substitution (SBS) signatures which revolutionised our ability to infer ongoing mutational mechanisms at the base-pair scale. However, current CIN signatures rely on correlations with mutations to determine driver mechanisms, potentially leading to inaccuracies. Moreover, they often reflect ancestral CIN mechanisms imprinted in the tumour genome detectable by bulk sequencing. Yet, CIN mechanisms may have evolved during tumourigenesis and can vary within the tumour or after treatment with genome-destabilizing therapies. Thus, decoding *ongoing* CIN mechanisms is crucial for effective therapy synergy.

Unfortunately, most available genomic alteration data comes from bulk sequencing of single end-point tumour samples from which detecting ongoing rates or types of CIN is not possible. While single-cell DNA sequencing is now emerging and reveals cell-cell heterogeneity in CNAs, it is still challenging to identify ongoing CIN-related copy-structural variations in static samples such as tumours, because determining which CNAs are recent or shaped by selection is ambiguous. Some progress has been made using computational approaches; a recent study measured the underlying tumour evolutionary processes to account for the effect of selection when quantitatively measuring CIN (Lynch et al. 2022). An alternative strategy to detect CNAs caused by chromosome mis-segregation events from large-scale single cell sequencing of cancer cell lines is to identify clones of cells based on similar CNA profiles. CNAs differing from the consensus within the clone are then assumed to be the consequence of recent CIN (Laks et al. 2019; Funnell et al. 2022). Previously, it has been demonstrated that whole chromosome missegregation events lead to whole copy-number alterations (CNAs) and that chromatin bridges can lead to sub chromosomal CNAs (Bolhaqueiro *et al*., 2019; Bollen *et al*., 2021). However, the contributing factor or deficit within the cell that results in these errors and CNAs is yet to be fully understood. Low numbers of cells, CNAs, and organoid samples precluded the detailed analysis of CNA features that could shed light on the causative mechanisms.

To determine in a controlled setting which CNAs were generated by ongoing cancer CIN, we devised a “Clone-Seq” workflow to track genomic evolution in single cancer cells as they form small populations by characterising newly-arising genomic alterations. Here, we track the evolution of two high grade serous ovarian cancer cell lines, which exhibit extreme rates of CIN. We show that both cell lines rapidly re-establish cell-cell heterogeneity to the level seen in the original parental population within 22 generations. We derive a workflow to extract the newly-arising copy number alterations from single cell genome sequencing data, and to time their occurrence to early, or late in the evolution of the population. We observe that some categories of CNA (chromosome arm-level losses and gains) appear to be under negative selection, and that others (focal amplifications) are more frequent in ancestral populations. In one cell line we observe one chromosome that is subject to a high rate of copy number alterations and functionally verify that this represents a chromosome undergoing extensive continual mis-segregation in mitosis.

## Results

### Single chromosomally unstable cancer cells re-establish genomic heterogeneity equivalent to parental populations within 22 generations

We chose two cell lines originating from high grade serous ovarian carcinoma (HGSOC); Kuramochi and OCM66. Kuramochi is a legacy cell line validated as being of HGSOC origin (Domcke *et al*., 2013) and previously characterised by our laboratory using functional genomics (Tamura *et al*., 2020b). OCM66 is one of a series of newly generated ovarian cancer models (OCMs) by the Taylor laboratory (Nelson *et al*., 2020). Both cell lines exhibit high rates of chromosome segregation errors during anaphase visible with microscopy (**Figure 1b-d**, and (Tamura *et al*., 2020b)). Typically, low density seeing, or FACS-based sorting of HGSOC cells results in a very poor viability. We therefore used the CellenONE microfluidics platform to seed single cells into individual wells of a 96 well plate, resulting in high efficiency of clone formation. We grew clones until they were near confluency in a 24 well plate, before sampling the final population using single cell DNA sequencing using either tagmentase-based (DLP+(Zahn *et al*., 2019)) or PCR-based(Bakker *et al*., 2016) workflows. We also sampled the originating (parental) populations. Parental populations and clones displayed visible cell-cell heterogeneity in terms of copy number profiles (**Figure 1e**). Despite this heterogeneity, pseudobulk analysis of the single cell data revealed that clones retained similar bulk karyotypic copy number alteration (CNA) profiles to parental populations (**Figure 1f, Figure S1a,b**). To quantify the cell-cell heterogeneity in the clonal populations we used two different metrics. First, we applied the phylogenetic tool MEDICC2, and calculated the average branch length – a measure of genomic evolutionary distance - between each individual final sampled cell and the inferred ancestor cell (**Figure 1g**). For both cell lines, cells from the newly derived clones exhibited equal evolutionary distance on average to those sampled from the parental populations, indicating that an equal level of genomic heterogeneity had been created by one single cell during clone growth. OCM66 cells showed a higher rate of diversification than Kuramochi (**Figure 1h**), suggesting higher rates of CIN. To examine diversity using an independent metric we devised a ‘dissimilarity score’ (see Methods). The cell-cell ‘dissimilarity score’ was also similar between clones and parental populations (**Figure 1i**). Together, these data demonstrate that high rates of CIN in both OCM66 and Kuramochi facilitate a rapid diversification in terms of copy number alterations from a single cell. Interestingly, this high rate of heterogeneity did not result in meaningful change in the bulk karyotype between clones and the parent population (**Figure 1f**), illustrating the importance of assessing the chromosomal alterations that occur ‘under the radar’, rather than bulk genomic alteration patterns. The interesting exception to this observation is an 11q gain which was lost from OCM66 clone 1 (**Figure 1f**), presumably due to the original seeding cell being one of the few in the parental population that did not carry this gain. Similarly in Kuramochi cells, we noted a copy number gain within chromosome 10q present in the parental population has been lost from Kuramochi clone 2, and a chromosome 5q monosomy lost from both clones (**Figure S1a,b**).

**Figure 1:**
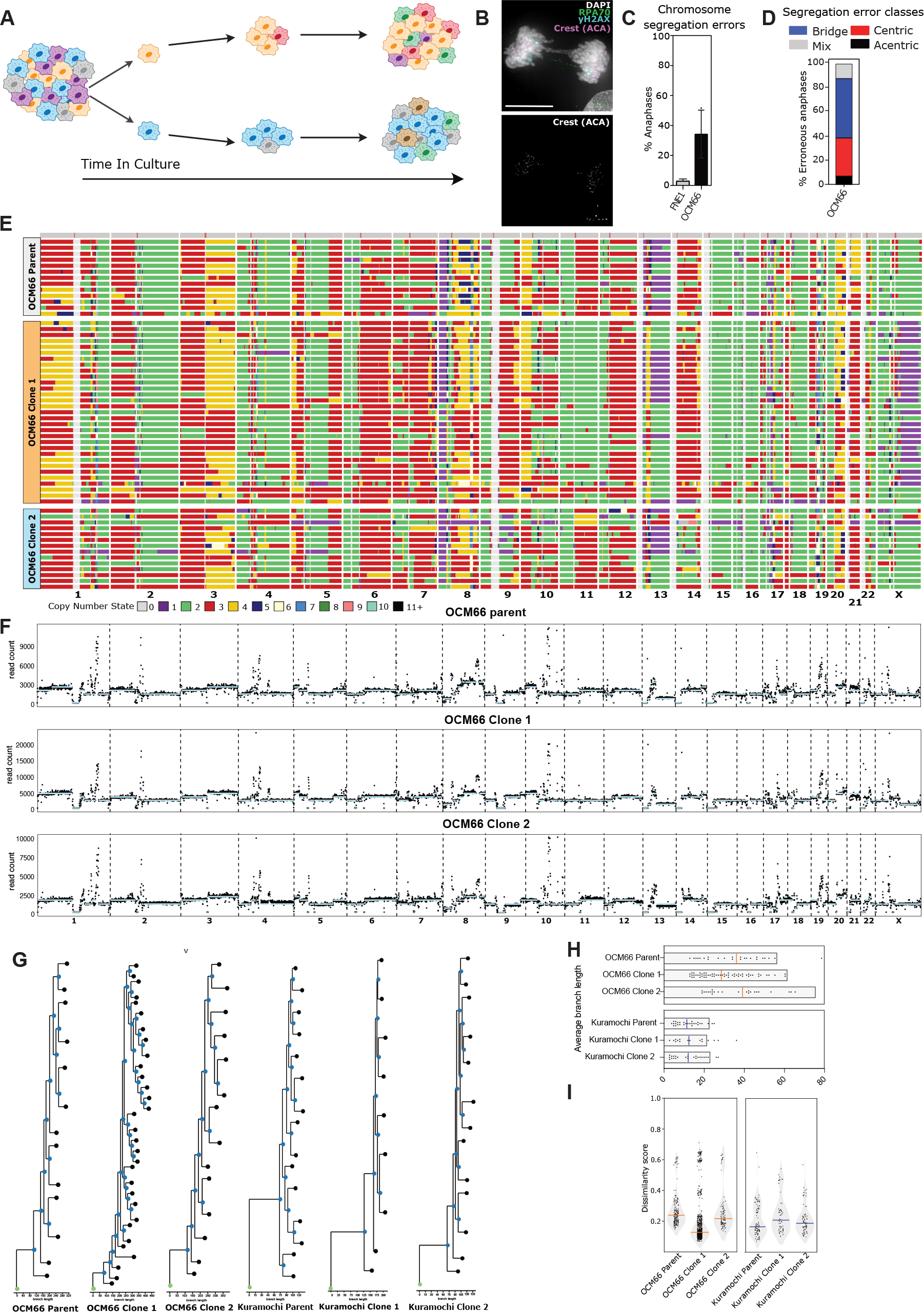
Single chromosomally unstable cancer cells re-establish genomic heterogeneity equivalent to parental populations within 22 generations. **A)** Schematic outlining the experimental strategy of single-cell outgrowth. **B)** Immunofluorescence images of OCM66 anaphase cells probed with antibodies to CREST (magenta; centromere) and RPA (green; ultra-fine bridge) and yH2AX (blue; DNA damage) and DAPI (grey; DNA) exhibiting a chromosome bridge and an ultrafine bridge. Scale bar: 5μm. **C)** OCM66 Chromosome segregation error rates quantified from fixed cell microscopy. **D)** Anaphase segregation errors from (C) classified according to error type. **E) L**ow-pass whole genome sequencing copy number heatmaps, each row represents a single cell, each column a chromosome and the colour (indicated by key) represents a different copy number. **F)** Pseudobulk copy number profiles, created by merging reads from all single cells to produce a bulk copy number profile. **G)** Copy-number based phylogenetic trees produced using MEDICC2 of HGSOC single cells from parental and clonal populations as indicated. **H)** Average branch lengths from analysis in (G). **I)** Dissimilarity score (see methods) calculated for each cell-cell pair in each indicated parent or clone population.

### Extracting the most recent copy number alterations from single cell clones highlights a specific unstable chromosome

We next focused on uncovering the most recent copy number alterations that had arisen during clonal outgrowth. Given that cancer cells typically deviate from diploidy, we first calculated each cell’s copy number profile relative to the inferred ancestral (the originating single cell of the clone) genome. To define the ancestral genome, we derived a reference genome comprised of the modal copy number state across binned regions of all cells in the clone, reasoning that alterations present in the originating cell would most often be retained as the most frequent (the mode) alteration in any subsequent daughter cell populations (**Figure 2a**). We then calculated all CNAs that deviated from the ancestral cell reference (**Figure 2b**). We noted that all cells within the clones accumulated new breakpoints at a similar rate during clonal outgrowth, implying a consistent rate of CIN among individual cells in the parental population, a measure that has been difficult to assess in previous studies **(Figure 2b)**.

**Figure 2:**
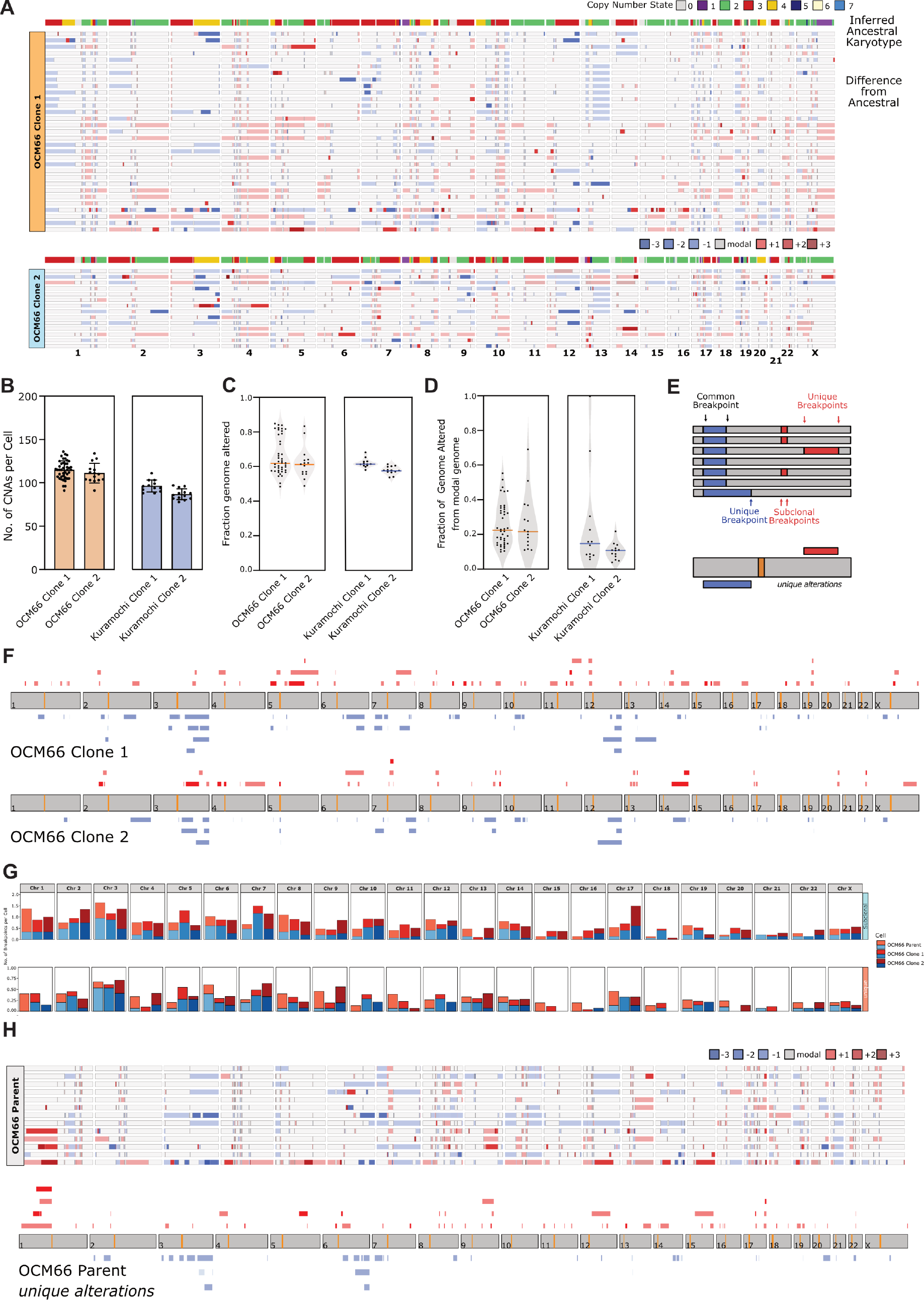
Extracting the most recent copy number alterations from single cell clones highlights a specific unstable chromosome. **A)** Copy number heatmaps for OCM66 clones 1 and 2, indicating the difference from inferred ancestral karyotype (modal) reference (see methods), with the corresponding ancestral reference above each heatmap. Heatmaps show the modal copy number in grey and copy number gains and losses in red-and blue-scale respectively according to the colour key. **B)** Total number of copy number alterations accumulated per cell that differ from the modal (ancestral) genome. **C)** Fraction of the genome altered (total length of altered regions/total genome length). **D)** Fraction of the genome altered after modal analysis (i.e. since clonal outgrowth). **E)** Schematic outlining the classification of unique CNAs, displaying a modal chromosome with CNAs that has shared and unique breakpoints. **F)** Diagram showing unique CNAs in each clonal populations after modal analysis and CNA filtering, red corresponds to CNA gains and blue to CNA losses, orange indicates the chromosome centromere. **G)** Graphical representation of the number of subclonal and unique CNAs per chromosome for each clonal population. **H)** Modal analysis (top) and unique CNAs (bottom) from parental population analysis.

We also calculated the fraction of genome altered (FGA), a frequently-used metric to infer CIN (Zhou et al. 2019) in two ways. First, we calculated FGA in the standard manner; assessing deviation from the normal (diploid) genome, which revealed that most of the genome is altered across all cells (**Figure 2c**), which is unsurprising given the highly rearranged nature of these genomes. Second, we calculated FGA *since clone outgrowth*, as an additional metric to assess the extent of genomic diversification during clone growth. Strikingly this analysis revealed that, during clonal outgrowth, OCM66 cells altered over 20% of their genomes on average, while Kuramochi cells altered ∼15% of their genomes (**Figure 2d**).

Next, we aimed to uncover those most recent CNAs, reasoning that these would most represent the alterations occurring due to ongoing CIN with the lowest interference from potential selection pressures. We achieved this by considering CNAs bounded by at least one unique breakpoint (among other cells in the clone) and occurring in only one cell within the clone population, and termed these ‘unique’ alterations (see scheme in **Figure 2e**). To visualize the frequency and distribution of these most recent CNAs across the genome, we created ‘pileups’ of all ‘unique’ alterations across a given clone plotted on a single ideogram (**Figure 2f; Figure S2b**). Intriguingly, in both OCM66 clones, we observed a high rate of recent alterations on chromosome 3 (**Figure 2f,g**), suggesting that this chromosome was unstable during clonal outgrowth of both clones, and therefore was likely to have been unstable in the parental population. To test this, we conducted a similar analysis of unique CNAs from the parental population, despite this population having been generated over much longer periods of time. Once again, chromosome 3 emerged as one of the most altered chromosomes in recent unique events (**Figure 2h**) though this could not be as clearly detected when considering all CNAs since clonal outgrowth (**Figure 2g**).

### Chromosome 3 in OCM66 cells is undergoing continual mis-segregation

We sought to investigate whether chromosome 3 indeed exhibited recurrent instability during cell division in OCM66 cells. To assess this, we performed Fluorescence In-Situ Hybridisation (FISH) on the parental OCM66 population using specific chromosome and centromere probes. We analysed the proportion of chromosome mis-segregation events that involved chromosome 3, comparing this to rates of mis-segregation of chromosomes 6 or 12 which serve as negative controls (low rates of CNAs; **Figure 2**). Strikingly, we observed that chromosome 3 was present in mis-segregating chromatin in over 35% of the cells analysed, most often in chromatin bridges, but also in lagging chromosome/chromosome fragments (**Figure 3a,b**). We then assessed the total contribution of this unstable chromosome to all mis-segregation events. Chromosome 3 accounted for almost one fifth of all chromosome segregation errors, compared to an expected 4.3% (expected frequency per chromosome). These data show that the clone-seq approach can identify specific chromosomes under particularly high rates of current instability, and demonstrate that cancer chromosomal instability can act in a highly biased nature towards specific chromosomes, as suggested previously (Shaikh *et al*., 2022a; Worrall *et al*., 2018; Klaasen *et al*., 2022).

**Figure 3:**
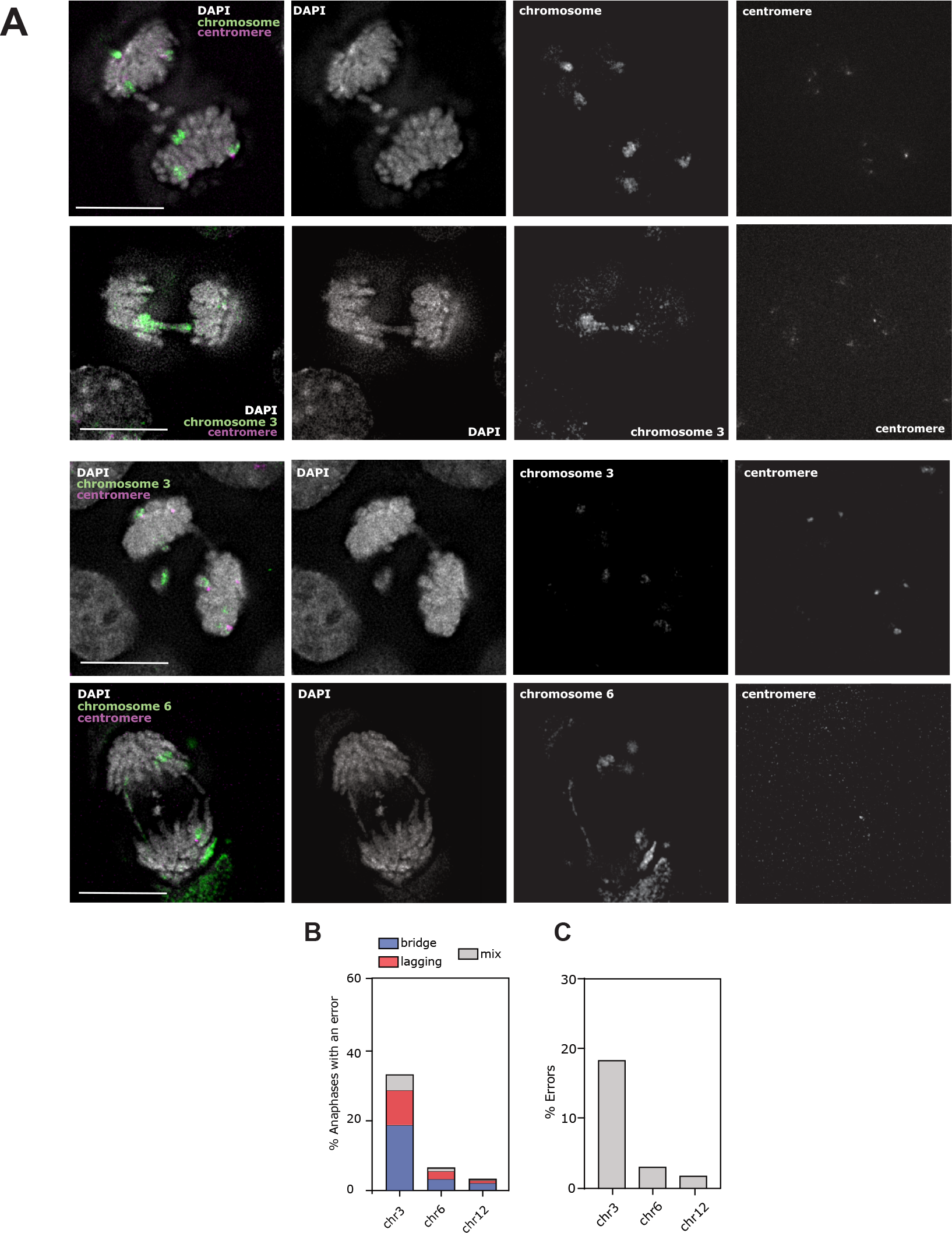
Chromosome 3 in OCM66 cells is under continual high rates of CIN. **A)** Representative microscopy images of OCM66 cells probed with fluorescence-in-situ hybridisation (FISH) probes for specific chromosomes and centromeres. Green: specific chromosome paint; grey: DAPI (DNA), magenta: chromosome specific centromere probe. Scale bar = 5μm. **B)** Quantification of the percentage of anaphase errors that are positive for chromosome indicated. **C)** Quantification of percentage of all errors that are positive for chromosome indicated.

### Analysis of most recent CNAs permits the detection of CNAs under negative selection

Since each cell we sequenced from clonal populations had undergone an average of 22 cell divisions, its genome serves as a comprehensive record of cumulative genomic evolution over that period. This allows us to explore potential differences in the spectrum of copy number alterations that occurred during the evolution from a single cell into a population. One particular question we sought to answer was whether we could identify evidence of negative selection against certain classes of CNAs. In traditional cancer genomics, measuring negative selection is challenging (one cannot observe events that are not there). We set out to compare the frequencies of specific CNA classes across different evolutionary timescales, ranging from early to mid-evolution and late evolution. We divided all the CNAs observed since clonal outgrowth into two categories; ‘subclonal’; any CNA observed since clonal outgrowth (excepting unique alterations), and ‘unique’; occurring in only one cell. We reasoned that alterations present in multiple cells are likely to have occurred earlier in clone evolution and thus are present in multiple cells. CNAs ranged in size, from focal (0.8 – 11.8 Mb) to chromosome arm-scale (>11.8 Mb – the length of the shortest chromosome arm (22p)), and size distributions were not different between subclonal or unique fractions (**Figure 4a,b**). We then analysed the proportions of focal vs chromosome-scale, and loss vs. gain, producing four CNA classes in combination. We performed a fold-change analysis of the proportion distributions between the subclonal, and unique CNAs for each clone (**Figure 4c,d**). Notably, unique CNAs from OCM66 clones exhibited an enrichment of chromosome-scale losses, while unique CNAs from Kuramochi clones displayed an enrichment of large gains, when compared to the subclonal alterations (**Figure 4c,d**). This suggests that chromosome-scale CNAs do occur at an appreciable rate, but are not maintained in the population, indicative of negative selection. We also observed that focal gains were depleted in unique CNAs; occurring less frequently than in the subclonal fraction. This suggests that focal gains may not occur very frequently, but are positively (or neutrally) selected when they do occur. Overall, these analyses suggest that it can be important to assess the CNAs occurring most recently in order to avoid selection effects.

**Figure 4:**
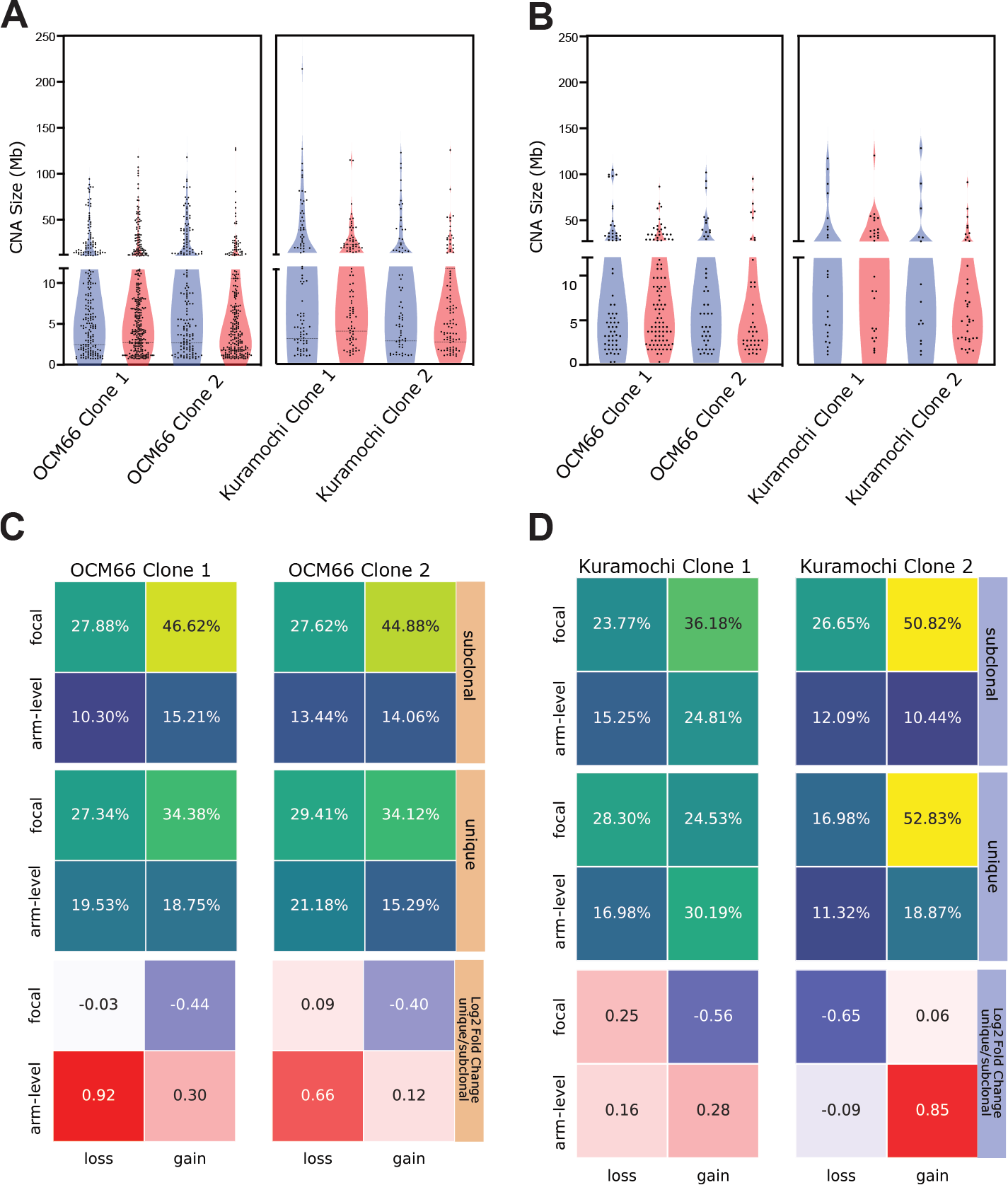
Analysis of most recent CNAs permits the detection of CNAs under negative selection. **A)** Scatter plot distribution of all CNA sizes that occurred since outgrowth split at 11.8 Mb (the smallest chromosome arm) **B)** Scatter plot of distribution of only unique CNA sizes, split at 11.8 Mb. **C-D)** Heatmaps showing the proportions of the types of CNA in each clonal population of OCM66 **(C)** and Kuramochi **(D)**. CNAs have been classified as subclonal (top) or unique (middle), and fold changes between unique and subclonal CNAs (bottom).

### Clonal outgrowth may provide more information than simply single cell sequencing parental populations

Our results above indicated that analysing the most recent CNAs from clonal outgrowth could be more informative of ongoing CIN-induced alterations. Such outgrowth experiments are not feasible in tumour samples however, so we sought to determine whether a similar approach in parental populations (analysis of the ‘unique’ CNAs) could be as informative. To test this, we performed a similar analysis as in Figure 4, but this time comparing the proportions of the four CNA classes in unique CNAs between parental, and clonal populations. We observed that the proportions differed substantially between parental, and clonal unique CNAs (**Figure 5a,b**), suggesting that the spectrum of CNAs extracted even from the ‘most recent’ events in parental populations may not fully capture all types of CNAs caused by ongoing CIN.

**Figure 5:**
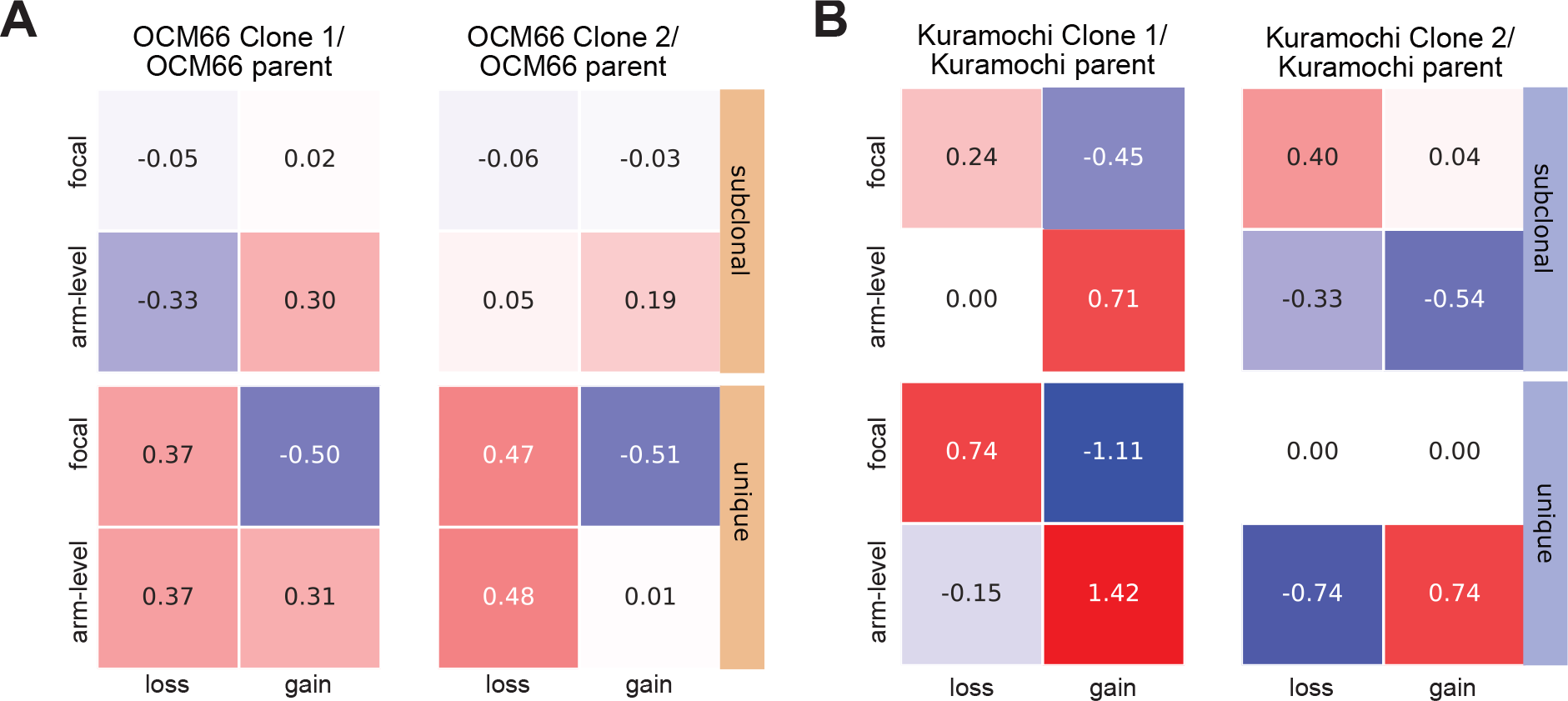
Clonal outgrowth may provide more information than simply single cell sequencing parental populations. **A,B)** Heatmaps showing fold changes between CNA sizes and types between parental populations and clones for OCM66 (A) and Kuramochi (B).

## Discussion

Here, we set out to design the optimal experimental strategy to define the genomic alterations caused by ongoing cancer CIN. Our ambition is to support and complement studies involving high throughput single cell sequencing of tumour samples and cancer cell lines, by rigorously testing whether sampling static populations can identify the full spectrum of CNAs generated by CIN.

In this study, we examined two cancer cell lines originating from high grade serous ovarian cancer, revealing these lines are capable of extremely rapid genomic re-diversification after only a few cell cycles. It will be very interesting to expand this study to additional cancer types. Previous experimental studies in colorectal cancer organoids did not detect such rapid re-diversification, perhaps due to a lower rate of CIN in these samples, or perhaps due to different resolution of single cell sequencing approaches used. To date, we have observed high rates of visible chromosome segregation errors in many different cancer types ((Tamura *et al*., 2020a; Burrell *et al*., 2013); our unpublished data), but it is likely that the majority of CNAs detected with single cell sequencing would be undetectable bymicroscopy, since we observed higher proportions of focal compared to chromosome-scale CNAs. Therefore, rates of CIN may not be accurately determined using microscopy alone.

We were able to detect frequent chromosome-scale alterations on chromosome 3 in OCM66 cells which correlated with this chromosome being consistently unstable during mitosis in the parental populations. Interestingly, these frequent alterations on chromosome 3 were mostly chromosome-scale deletions, suggesting loss of chromosomal material perhaps through loss of genetic material from the nucleus, such as into micronuclei.

When we compared the proportions of CNAs between subclonal and unique CNAs, we noted that across both cell lines and all clones, chromosome-scale alterations appeared to be under negative selection. This is perhaps not surprising since many of the large deletions are associated with absolute (not relative to modal karyotype) copy number of 1 (monosomy), which is very poorly tolerated in non-transformed cells (Chunduri et al., 2021). Paradoxically however, recurrent monosomy is observed in several chromosome regions in these cell lines. For example, chromosomes 8p and 13q in OCM66 cells, and Xq in OCM66 clone 1 (**Figure 1b**)), and chromosomes 5q, 18 and X in Kuramochi (Figure S1a). This suggests that spontaneous large deletions are generally detrimental, even to cancer cells with highly rearranged genomes, unless they are associated with a fitness benefit. It will be important to delve into the potential reasons why these specific chromosomal regions are tolerated, given the overall negative fitness of large deletions in general.

Although our study suggests that recent outgrowth of cancer cells provides the most accurate picture of CIN-induced genome evolution, we believe that these findings can be integrated into analysis of cancer sequencing from static samples. For example, building a complete picture of which alterations tend to be under negative selection can help the interpretation of CNAs from single cell sequencing of tumours.

## Methods

### Cell Lines

Kuramochi cells were purchased from the Japanese Collection of Research Bioresources and maintained in RPMI (Gibco) supplemented with 10% FBS and 100 U penicillin/streptomycin. OCM66 was a kind gift from Prof Stephen Taylor (University of Manchester). OCM66 was maintained in OCMI (Ince et al., 2015). OCMI is made using a 50:50 mix of Nutrient Mixture Ham’s F12 (Sigma Aldrich) and Medium 199 (Life Technologies) supplemented with 5% Hyclone FBS (GE Healthcare), 0.5 ng/ml 17 β-oestradiol, 5 μg/ml all-trans retinoic acid, 0.012 μg/ml ascorbic acid, 0.003 μg/ml α-tocopherol phosphate, 1.25 mg/ml BSA, 0.025 μg/ml calciferol, 25 ng/ml cholera toxin, 0.05 μg/ml cholesterol, 3.5 μg/ml choline chloride, 0.01 μg/ml EGF; 2 mM glutamine (Sigma Aldrich), 0.33 μg/ml folic acid, 10 mM HEPES at pH7.4, 0.5 μg/ml hydrocortisone, 1.75 μg/ml hypoxanthine, 4.5 μg/ml i-inositol, 20 μg/ml insulin, 0.05 μg/ml lipoic acid, 5 μg/ml o-phosphoryl ethanolamine, 0.0125 μg/ml para-aminobenzioic acid, 100 U/ml penicillin/streptomycin (Sigma Aldrich), 0.125 μg/ml ribose, 8 ng/ml selenious acid, 0.08 μg/ml thiamine HCL, 10 μg/ml transferrin, 0.2 pg/ml Tridothyronine, 0.075 μg/ml uracil, 0.35 μg/ml vitamin B12, 0.085 μg/ml xanthine (all from Sigma). Cells are routinely tested for presence of mycoplasma using MycoAlert PLUS Mycoplasma Detection Kit (LT07-710, Lonza).

### Immunofluorescence

OCM66 cells were grown on coverslips coated with collagen and fixed with PTEMF (PIPES (pH 6.8), 0.2% Triton, 0.01M EGTA, 1mmol/L MgCl_2_, 4% Formaldehyde). Coverslips were blocked with 3% BSA and incubated with primary antibodies (RPA - ab79398, H2AX - Millipore-05-636, CREST - Antibodies incorporated - 15-234-0001). Then secondary antibodies (goat anti-rabbit AlexaFluor (AF)488, goat anti-mouse AF594 (Invitrogen - A11008, A11001) and goat anti-human AF647 (Stratech 109-606-088- JIR)) DNA was then stained with DAPI (Roche) and coverslips mounted with Vectashield (Vector Laboratories - Vector H-1000).

### Chromosome Painting

OCM66 cells were grown on microscope slides coated with collagen and fixed with 3:1 Methanol:Acetic Acid then subject to sequential ethanol dehydration (2 minutes in 70%, 90% then 100% ethanol) then air dried. Chromosome and centromere probes (Cytocell) were added to slides and then heated to 72° for 2 minutes, then left at 37° in a humid chamber overnight. Slides were then washed in 0.25 SSC at 72° for 2 minutes then in 2xSSC, 0.01% Tween at room temperature for 30 seconds. Slides were then stained with DAPI and coverslips mounted with Vectashield.

## Microscopy

Images were acquired using an Olympus DeltaVision RT microscope (Applied Precision, LLC) equipped with a CoolSnap HQ camera. 3D image stacks were acquired at 0.2μm intervals, using an Olympus 100x 1.4 numerical aperture UPlanSApo oil immersion objective. Deconvolution and analysis was performed using SoftWorxExplorer (Applied Precision, LLC).

### Single cell clonal outgrowth and sequencing

Single cells were dispensed into 96 well plates using the CellenOne microfluidics system. Clonal populations were monitored until confluence or overlapping growth and moved to a 24-well plate until confluence at which time cells were counted and stored in freezing medium until sequencing. The number of cell generations that occurred since the starting cell (*n*) was estimated using the relationship 2^*n*^ = final cell count.

Single cells were dispensed into 384 Lo-bind DNA plates containing unique pairs of Nextera i5 and i7 primers using CellenOne microfluidics. OCM66 library preparation followed the tagmentase based DLP+ method described previously (Zahn *et al*., 2019). Demultiplexing was performed based on unique-barcode identifiers using bcl2fastq (v1.8.4, Illumina). Demultiplexed reads were trimmed using trimmomatic using Nextera adaptors and length quality parameters and QC was performed using fastqc. Trimmed reads were aligned to GRCh38 genomic reference using bwa-mem, only reads over a map quality of 10 are used for downstream analysis. For Kuramochi, library preparation used a PCR based method where single nuclei were isolated and stained with 10 μg/mL propidium iodide and 10 μg/mL Hoechst was used. Single nuclei with low Hoechst/PI fluorescence (G1 population) were sorted into 96-well plates containing freezing buffer using a FACSJazz (BD Biosciences). Pre-amplification-free single-cell whole genome sequencing libraries were prepared using a Bravo Automated Liquid Handling Platform (Agilent Technologies, Santa Clara, CA, USA), followed by size-selection and extraction from a 2% E-gel EX (Invitrogen). Single-end 84-nt sequence reads were generated using the NextSeq 500 system (Illumina, San Diego, CA, USA) at 192 single-cell DNA libraries per flow cell. Demultiplexing based on library-specific barcodes and conversion to fastq format was done using bcl2fastq (v1.8.4, Illumina). Duplicate reads were called using BamUtil (v1.0.3). Demultiplexed reads were aligned to the GRCh38 reference genome using bowtie (v2.2.4), and only uniquely mapped reads (MAPQ>10) were used for further analysis. To validate the comparison of the two sequencing methods we utilised previous data produced by the Taylor lab (Nelson *et al*., 2020) using the PCR-based method of OCM66 single cells, with our DLP+ data. Here we utilised the dissimilarity score to confirm there were no major dissimilarities or artifacts from the sequencing methods (**Figure S3**). Copy number analysis was performed using AneuFinder (v1.26.0) using 500kb bins, GC correction and blacklisting, then edivisive was used to determine the most likely copy number states. We only analysed cells between 0.5M-3M reads/cell, and verified even coverage along the genome in order to ensure an equal ability to detect CNAs across all cells and discarded any CNAs below 0.8 Mb for further analysis. Each sample was also analysed using the developer version of AneuFinder (GitHub) where the ground ploidy was constrained between 2 and 3. Afterwards each sample was subject to modal normalisation, in 500kb sliding windows the modal copy number was calculated, creating a reference used to re-analyse each cell. Any data that had less than 20 reads per bin were discarded.

### Heterogeneity and aneuploidy scoring

To measure the unlikeness and distance between cell CNA profiles, we developed a dissimilarity score based on the combination of two similarity measures: cosine similarity and euclidean distance. Cosine similarity measures the cosine of the angle between two vectors and ranges from -1 to 1, where 1 indicates identical directions, 0 indicates orthogonality (no similarity), and -1 indicates opposite directions. Cosine dissimilarities are measured by subtracting 1 from cosine similarities. The Euclidean distance measures the straight-line distance between two points in Euclidean space. It considers the magnitude of the vectors and provides a non-negative value. To have measures within the range of 0 to 1, distances were max-normalized. Therefore, the resulting dissimilarity score ranges between 0 and 1, where 0 indicates maximum similarity, and 1 indicates maximum dissimilarity.

Let ***Sc*** represent the cosine similarity between two vectors, and ***Se*** represent the normalized Euclidean distance between the same vectors. The dissimilarity score (***D***) can be defined as a function of ***Sc*** and ***Se*** as follows:

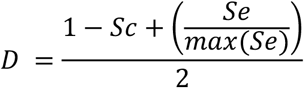

To calculate dissimilarity scores, genomes were divided in sliding windows of 500Kb with a step size of 1/3 the window size, having a total of 3 windows per 500Kb. The sliding windows were intersected with CNA profiles to calculate the mean total CN of each window. The dissimilarity scores were calculated from these vectors.

### Subsetting of clonal, subclonal and unique CNAs

Any CNA removed by modal analysis was considered clonal. Those that remained were further filtered. A CNA was considered to be unique if at least one of the breakpoint locations did not overlap with any other CNA breakpoint, allowing for 1 Mb error. Any ‘unique’ breakpoints that occurred at either end of the chromosome were ignored to prevent mapping artefacts. CNA filtering and measurement scripts are available at https://github.com/MBoemo/clonalMasker and a variant of clonalMasker which filters via breakpoint rather than whole CNAs.

## Acknowledgements

We acknowledge the kind gift of OCM66 from Professor Stephen Taylor.

## Author contributions

MG performed all cell biological and most computational analyses and wrote the paper, supervised by SM. DG-P performed computational analysis supervised by SM and CS. AA and CVH performed computational analyses, supervised by SM. TS performed cell biology experiments, supervised by SM. BB, DCJS and RW performed PCR-based single cell sequencing and computational analyses, supervised by FF. RS assisted with MEDICC2 implementation and analysis. SM conceived the study, designed the experiments, analysed the data and wrote the paper.

## Declarations

The authors declare no conflicts of interest.

## Supplementary Figures

**Supplementary Figure 1.**
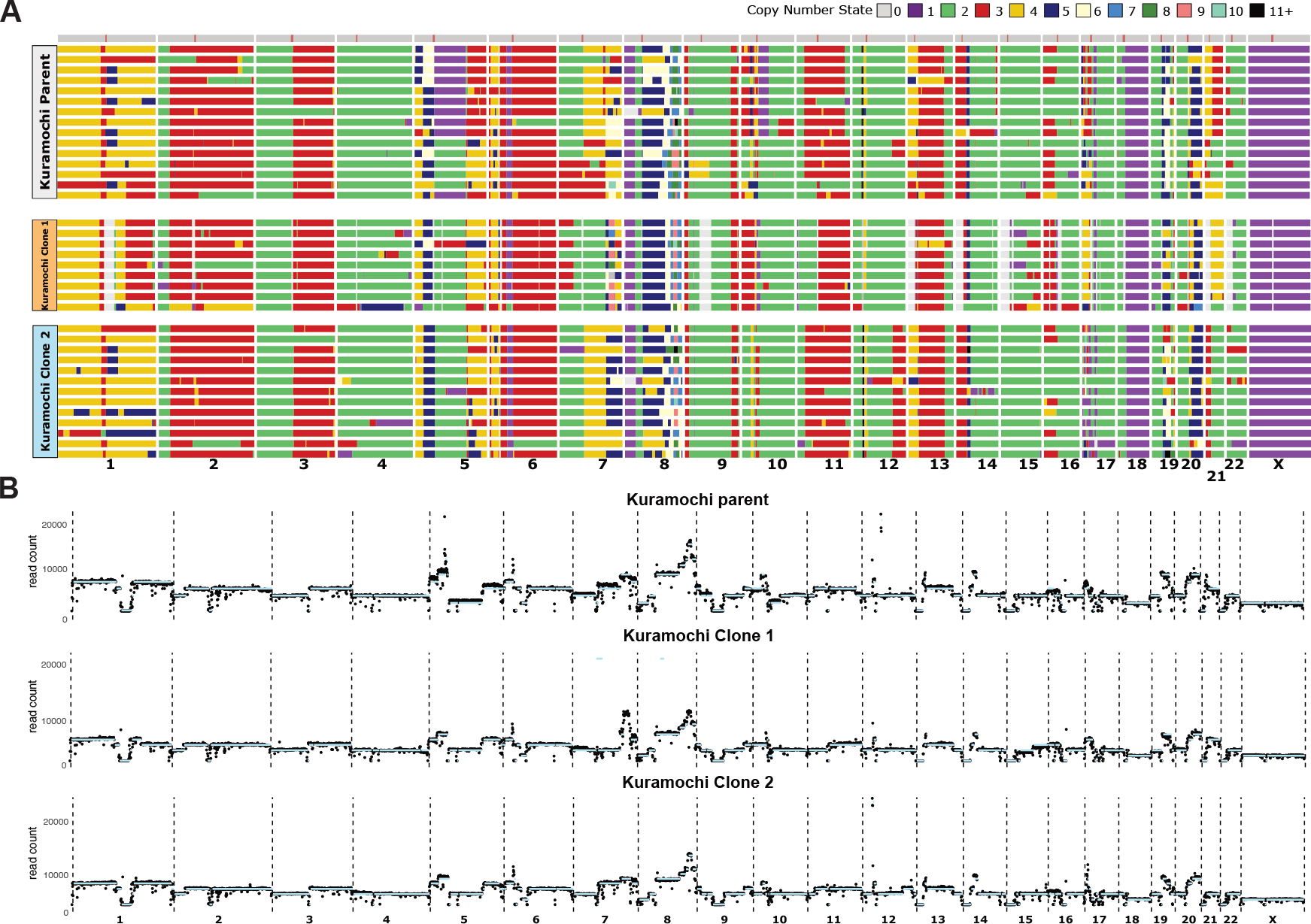
**A) L**ow-pass whole genome sequencing copy number heatmaps, each row represents a single cell, each column a chromosome and the colour (indicated by key) represents a different copy number. **B)** Pseudobulk copy number profiles, created by merging reads from all single cells to produce a bulk copy number profile.

**Supplementary Figure 2.**
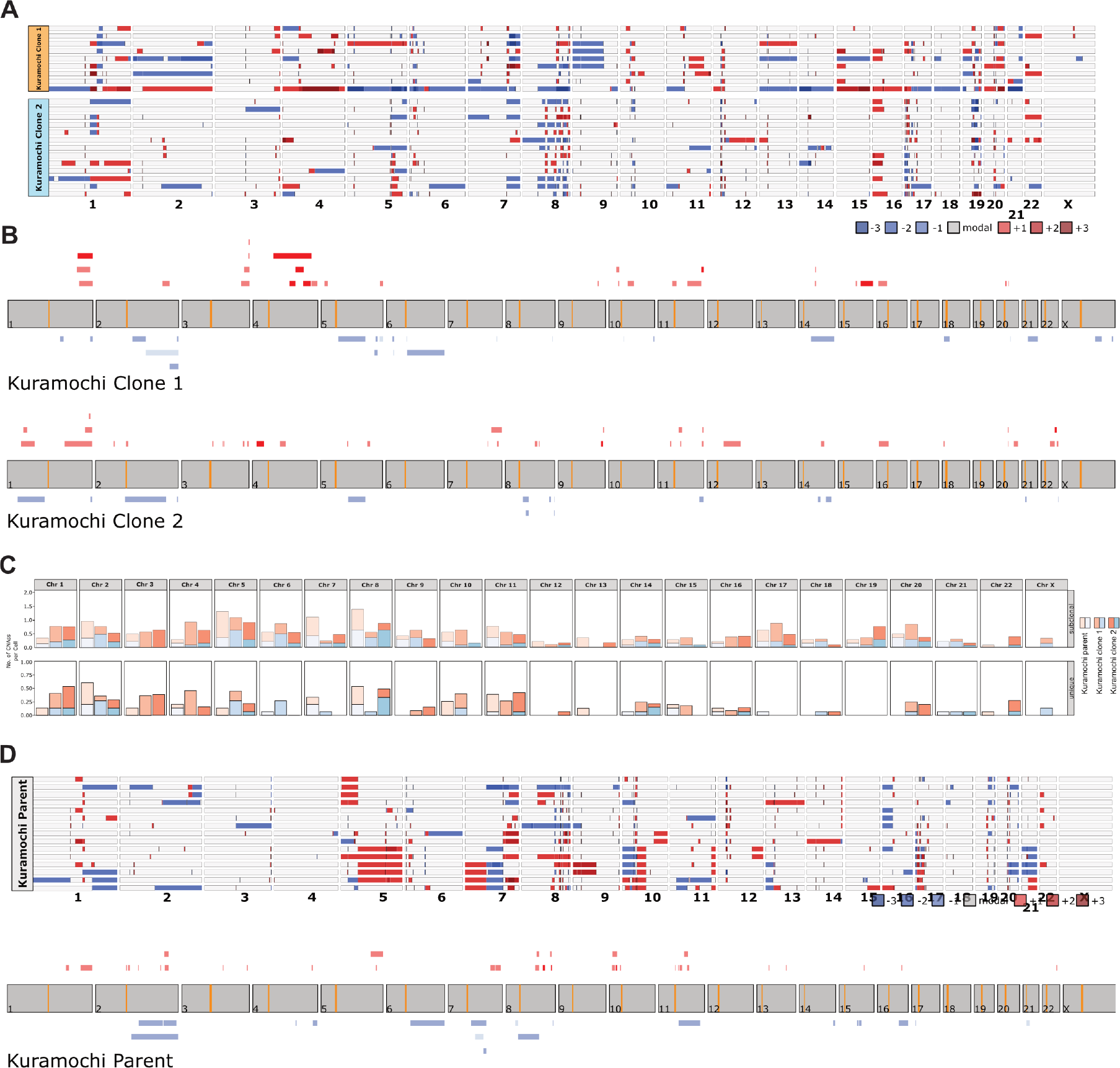
**A)** Copy number heatmaps indicating the difference from inferred ancestral karyotype (modal) reference (see methods), with the corresponding ancestral reference above each heatmap. Heatmaps show the modal copy number in grey and copy number gains and losses in reds and blues respectively according to the colour key. **B)** Diagram showing unique CNAs in Kuramochi clonal populations after modal analysis and CNA filtering, red corresponds to CNA gains and blue to CNA losses, orange indicates the chromosome centromere **C)** Graphical representation of the number of subclonal and unique CNAs per chromosome for each clonal population. **D)** Modal analysis (top) and unique CNAs (bottom) from parental population analysis.

**Supplementary Figure 3.**
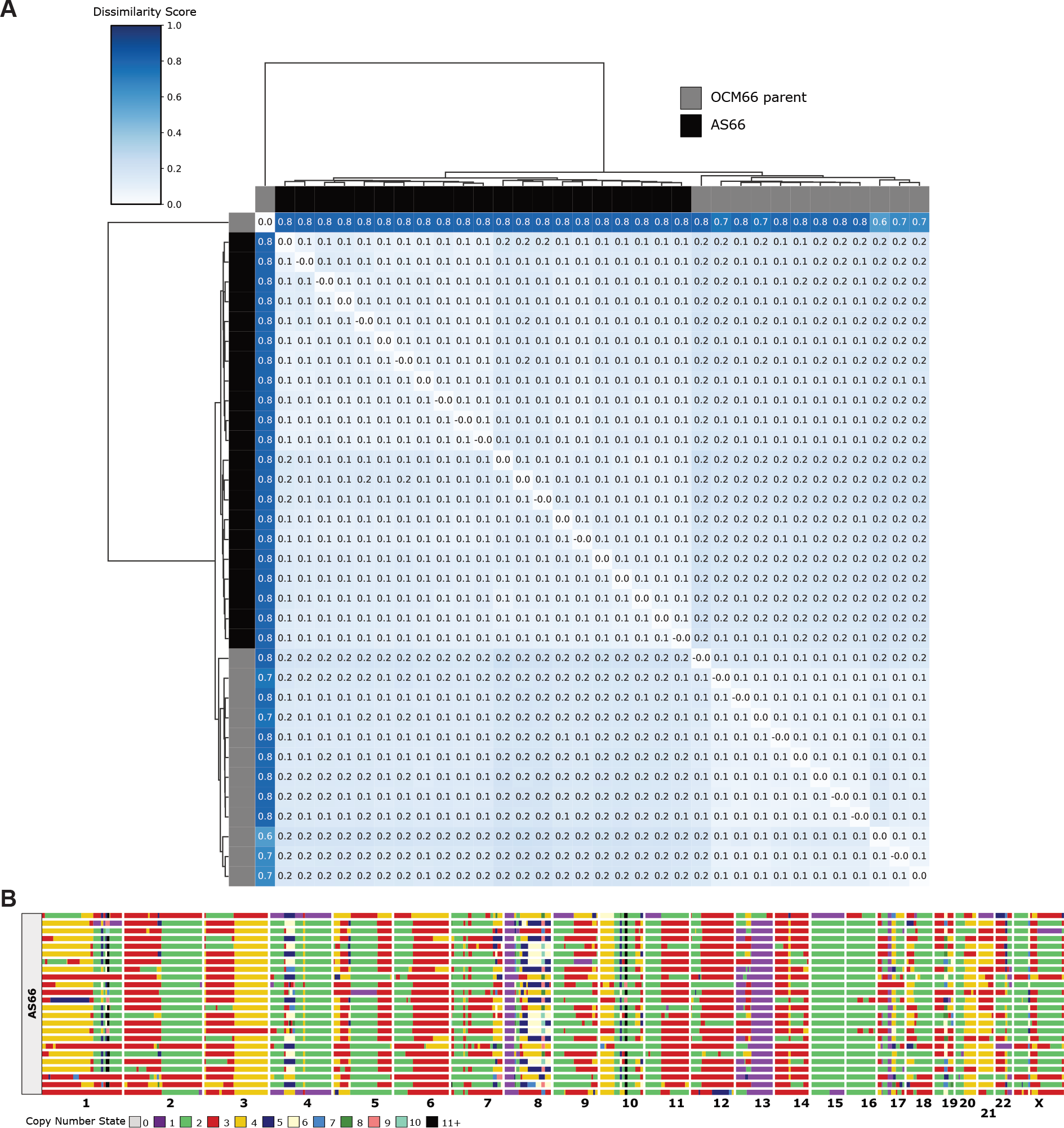
**A)** Dissimilarity heatmap analysis of OCM66 parental population and original scWGS performed in (Nelson *et al*., 2020). **B)** Replotted copy number heatmap of original scWGS data from (Nelson *et al*., 2020).

## References

Bakhoum, S. F., Genovese, G. and Compton, D. A. (2009) ‘Deviant kinetochore microtubule dynamics underlie chromosomal instability’, Curr Biol, 19(22), pp. 1937–42.

Bakker, B., Taudt, A., Belderbos, M. E., Porubsky, D., Spierings, D. C., de Jong, T. V., Halsema, N., Kazemier, H. G., Hoekstra-Wakker, K., Bradley, A., de Bont, E. S., van den Berg, A., Guryev, V., Lansdorp, P. M., Colome-Tatche, M. and Foijer, F. (2016) ‘Single-cell sequencing reveals karyotype heterogeneity in murine and human malignancies’, Genome Biol, 17(1), pp. 115.

Bolhaqueiro, A. C. F., Ponsioen, B., Bakker, B., Klaasen, S. J., Kucukkose, E., van Jaarsveld, R. H., Vivié, J., Verlaan-Klink, I., Hami, N., Spierings, D. C. J., Sasaki, N., Dutta, D., Boj, S. F., Vries, R. G. J., Lansdorp, P. M., van de Wetering, M., van Oudenaarden, A., Clevers, H., Kranenburg, O., Foijer, F., Snippert, H. J. G. and Kops, G. J. P. L. (2019) ‘Ongoing chromosomal instability and karyotype evolution in human colorectal cancer organoids’, Nat Genet, 51(5), pp. 824–834.

Bollen, Y., Stelloo, E., van Leenen, P., van den Bos, M., Ponsioen, B., Lu, B., van Roosmalen, M. J., Bolhaqueiro, A. C. F., Kimberley, C., Mossner, M., Cross, W. C. H., Besselink, N. J. M., van der Roest, B., Boymans, S., Oost, K. C., de Vries, S. G., Rehmann, H., Cuppen, E., Lens, S. M. A., Kops, G. J. P. L., Kloosterman, W. P., Terstappen, L. W. M. M., Barnes, C. P., Sottoriva, A., Graham, T. A. and Snippert, H. J. G. (2021) ‘Reconstructing single-cell karyotype alterations in colorectal cancer identifies punctuated and gradual diversification patterns’, Nat Genet, 53(8), pp. 1187–1195.

Bryant, H. E., Schultz, N., Thomas, H. D., Parker, K. M., Flower, D., Lopez, E., Kyle, S., Meuth, M., Curtin, N. J. and Helleday, T. (2005) ‘Specific killing of BRCA2-deficient tumours with inhibitors of poly(ADP-ribose) polymerase.’, Nature, 434(7035), pp. 913–917.

Burrell, R. A., McClelland, S. E., Endesfelder, D., Groth, P., Weller, M. C., Shaikh, N., Domingo, E., Kanu, N., Dewhurst, S. M., Gronroos, E., Chew, S. K., Rowan, A. J., Schenk, A., Sheffer, M., Howell, M., Kschischo, M., Behrens, A., Helleday, T., Bartek, J., Tomlinson, I. P. and Swanton, C. (2013) ‘Replication stress links structural and numerical cancer chromosomal instability’, Nature, 494(7438), pp. 492–6.

Chunduri, N. K., Menges, P., Zhang, X., Wieland, A., Gotsmann, V. L., Mardin, B. R., Buccitelli, C., Korbel, J. O., Willmund, F., Kschischo, M., Raeschle, M. and Storchova, Z. (2021) ‘Systems approaches identify the consequences of monosomy in somatic human cells’, Nat Commun, 12(1), pp. 5576.

Davies, H., Glodzik, D., Morganella, S., Yates, L. R., Staaf, J., Zou, X., Ramakrishna, M., Martin, S., Boyault, S., Sieuwerts, A. M., Simpson, P. T., King, T. A., Raine, K., Eyfjord, J. E., Kong, G., Borg, A., Birney, E., Stunnenberg, H. G., van de Vijver, M. J., Borresen-Dale, A. L., Martens, J. W., Span, P. N., Lakhani, S. R., Vincent-Salomon, A., Sotiriou, C., Tutt, A., Thompson, A. M., Van Laere, S., Richardson, A. L., Viari, A., Campbell, P. J., Stratton, M. R. and Nik-Zainal, S. (2017) ‘HRDetect is a predictor of BRCA1 and BRCA2 deficiency based on mutational signatures’, Nat Med, 23(4), pp. 517–525.

Domcke, S., Sinha, R., Levine, D. A., Sander, C. and Schultz, N. (2013) ‘Evaluating cell lines as tumour models by comparison of genomic profiles’, Nat Commun, 4, pp. 2126.

Ertych, N., Stolz, A., Stenzinger, A., Weichert, W., Kaulfuß, S., Burfeind, P., Aigner, A., Wordeman, L. and Bastians, H. (2014) ‘Increased microtubule assembly rates influence chromosomal instability in colorectal cancer cells’, Nature cell biology.

Farmer, H., McCabe, N., Lord, C. J., Tutt, A. N. J., Johnson, D. A., Richardson, T. B., Santarosa, M., Dillon, K. J., Hickson, I., Knights, C., Martin, N. M. B., Jackson, S. P., Smith, G. C. M. and Ashworth, A. (2005) ‘Targeting the DNA repair defect in BRCA mutant cells as a therapeutic strategy.’, Nature, 434(7035), pp. 917–921.

Forment, J. V. and O’Connor, M. J. (2018) ‘Targeting the replication stress response in cancer’,Pharmacol Ther, 188, pp. 155–167.

Klaasen, S. J., Truong, M. A., van Jaarsveld, R. H., Koprivec, I., Stimac, V., de Vries, S. G., Risteski, P., Kodba, S., Vukusic, K., de Luca, K. L., Marques, J. F., Gerrits, E. M., Bakker, B., Foijer, F., Kind, J., Tolic, I. M., Lens, S. M. A. and Kops, G. (2022) ‘Nuclear chromosome locations dictate segregation error frequencies’, Nature, 607(7919), pp. 604–609.

Macintyre, G., Goranova, T. E., De Silva, D., Ennis, D., Piskorz, A. M., Eldridge, M., Sie, D., Lewsley, L. A., Hanif, A., Wilson, C., Dowson, S., Glasspool, R. M., Lockley, M., Brockbank, E., Montes, A., Walther, A., Sundar, S., Edmondson, R., Hall, G. D., Clamp, A., Gourley, C., Hall, M., Fotopoulou, C., Gabra, H., Paul, J., Supernat, A., Millan, D., Hoyle, A., Bryson, G., Nourse, C., Mincarelli, L., Sanchez, L. N., Ylstra, B., Jimenez-Linan, M., Moore, L., Hofmann, O., Markowetz, F., McNeish, I. A. and Brenton, J. D. (2018) ‘Copy number signatures and mutational processes in ovarian carcinoma’, Nat Genet, 50(9), pp. 1262–1270.

McBride, D. J., Etemadmoghadam, D., Cooke, S. L., Alsop, K., George, J., Butler, A., Cho, J., Galappaththige, D., Greenman, C., Howarth, K. D., Lau, K. W., Ng, C. K., Raine, K., Teague, J., Wedge, D. C., Cancer Study Group, A. O., Caubit, X., Stratton, M. R., Brenton, J. D., Campbell, P. J., Futreal, P. A. and Bowtell, D. D. (2012) ‘Tandem duplication of chromosomal segments is common in ovarian and breast cancer genomes’, J Pathol, 227(4), pp. 446–55.

Nelson, L., Tighe, A., Golder, A., Littler, S., Bakker, B., Moralli, D., Baker, S. M., Donaldson, I. J., Spierings, D. C. J., Wardenaar, R., Neale, B., Burghel, G. J., Winter-Roach, B., Edmondson, R., Clamp, A. R., Jayson, G. C., Desai, S., Green, C. M., Hayes, A., Foijer, F., Morgan, R. D. and Taylor, S. S. (2020) ‘A living biobank of ovarian cancer ex vivo models reveals profound mitotic heterogeneity’, Nature communications, 11(1), pp. 1 –18.

Ng, C. K., Cooke, S. L., Howe, K., Newman, S., Xian, J., Temple, J., Batty, E. M., Pole, J. C., Langdon, S. P., Edwards, P. A. and Brenton, J. D. (2012) ‘The role of tandem duplicator phenotype in tumour evolution in high-grade serous ovarian cancer’, J Pathol, 226(5), pp. 703–12.

Nik-Zainal, S., Davies, H., Staaf, J., Ramakrishna, M., Glodzik, D., Zou, X., Martincorena, I., Alexandrov, L. B., Martin, S., Wedge, D. C., Van Loo, P., Ju, Y. S., Smid, M., Brinkman, A. B., Morganella, S., Aure, M. R., Lingjaerde, O. C., Langerod, A., Ringner, M., Ahn, S. M., Boyault, S., Brock, J. E., Broeks, A., Butler, A., Desmedt, C., Dirix, L., Dronov, S., Fatima, A., Foekens, J. A., Gerstung, M., Hooijer, G. K., Jang, S. J., Jones, D. R., Kim, H. Y., King, T. A., Krishnamurthy, S., Lee, H. J., Lee, J. Y., Li, Y., McLaren, S., Menzies, A., Mustonen, V., O’Meara, S., Pauporte, I., Pivot, X., Purdie, C. A., Raine, K., Ramakrishnan, K., Rodriguez-Gonzalez, F. G., Romieu, G., Sieuwerts, A. M., Simpson, P. T., Shepherd, R., Stebbings, L., Stefansson, O. A., Teague, J., Tommasi, S., Treilleux, I., Van den Eynden, G. G., Vermeulen, P., Vincent-Salomon, A., Yates, L., Caldas, C., van’t Veer, L., Tutt, A., Knappskog, S., Tan, B. K., Jonkers, J., Borg, A., Ueno, N. T., Sotiriou, C., Viari, A., Futreal, P. A., Campbell, P. J., Span, P. N., Van Laere, S., Lakhani, S. R., Eyfjord, J. E., Thompson, A. M., Birney, E., Stunnenberg, H. G., van de Vijver, M. J., Martens, J. W., Borresen-Dale, A. L., Richardson, A. L., Kong, G., Thomas, G. and Stratton, M. R. (2016) ‘Landscape of somatic mutations in 560 breast cancer whole-genome sequences’, Nature, 534(7605), pp. 47–54.

Shaikh, N., Mazzagatti, A., Angelis, S. D., Johnson, S. C., Bakker, B., Spierings, D. C. J., Wardenaar, R., Maniati, E., Wang, J., Boemo, M. A., Foijer, F. and McClelland, S. E. (2022a) ‘Replication stress generates distinctive landscapes of DNA copy number alterations and chromosome scale losses’, Genome Biology, pp. 1 –24.

Shaikh, N., Mazzagatti, A., De Angelis, S., Johnson, S. C., Bakker, B., Spierings, D. C. J., Wardenaar, R., Maniati, E., Wang, J., Boemo, M. A., Foijer, F. and McClelland, S. E. (2022b) ‘Replication stress generates distinctive landscapes of DNA copy number alterations and chromosome scale losses’, Genome Biol, 23(1), pp. 223.

Tamura, N., Shaikh, N., Muliaditan, D., Soliman, T. N., McGuinness, J. R., Maniati, E., Moralli, D., Durin, M.-A., Green, C. M., Balkwill, F. R., Wang, J., Curtius, K. and McClelland, S. E. (2020a) ‘Specific Mechanisms of Chromosomal Instability Indicate Therapeutic Sensitivities in High-Grade Serous Ovarian Carcinoma’, Cancer research, 80(22), pp. 4946 –4959.

Tamura, N., Shaikh, N., Muliaditan, D., Soliman, T. N., McGuinness, J. R., Maniati, E., Moralli, D., Durin, M. A., Green, C. M., Balkwill, F. R., Wang, J., Curtius, K. and McClelland, S. E. (2020b) ‘Specific Mechanisms of Chromosomal Instability Indicate Therapeutic Sensitivities in High-Grade Serous Ovarian Carcinoma’, Cancer Res, 80(22), pp. 4946–4959.

Thompson, S. L. and Compton, D. A. (2011) ‘Chromosome missegregation in human cells arises through specific types of kinetochore-microtubule attachment errors’, Proceedings of the National Academy of Sciences, 108(44), pp. 17974 –17978.

Ubhi, T. and Brown, G. W. (2019) ‘Exploiting DNA Replication Stress for Cancer Treatment’, Cancer Res, 79(8), pp. 1730–1739.

Worrall, J. T., Tamura, N., Mazzagatti, A., Shaikh, N., van Lingen, T., Bakker, B., Spierings, D. C. J., Vladimirou, E., Foijer, F. and McClelland, S. E. (2018) ‘Non-random Mis-segregation of Human Chromosomes’, Cell Reports, 23(11), pp. 3366–3380.

Zahn, H. and Lai, D. and Steif, A. and Brimhall, J. and Biele, J. and Wang, B. and Masud, T. and Ting, J. and Grewal, D. and Nielsen, C. and Leung, S. and Bojilova, V. and Smith, M. and Golovko, O. and Poon, S. and Eirew, P. and Kabeer, F. and Algara, T. R. d. and Lee, S. R. and Taghiyar, M. J. and Huebner, C. and Ngo, J. and Chan, T. and Vatrt-Watts, S. and Walters, P. and Abrar, N. and Chan, S. and Wiens, M. and Martin, L. and Scott, R. W. and Underhill, T. M. and Chavez, E. and Steidl, C. and Costa, D. D. and Ma, Y. and Coope, R. J. N. and Corbett, R. and Pleasance, S. and Moore, R. and Mungall, A. J. and Mar, C. and Cafferty, F. and Gelmon, K. and Chia, S. and Team, T. C. I. G. C. and Hannon, G. J. and Battistoni, G. and Bressan, D. and Cannell, I. and Casbolt, H. and Jauset, C. and Kovačević, T. and Mulvey, C. and Nugent, F. and Ribes, M. P. and Pearsall, I. and Qosaj, F. and Sawicka, K. and Wild, S. and Williams, E. and Aparicio, S. and Laks, E. and Li, Y. and O’Flanagan, C. and Smith, A. and Ruiz, T. and Balasubramanian, S. and Lee, M. and Bodenmiller, B. and Burger, M. and Kuett, L. and Tietscher, S. and Windager, J. and Boyden, E. and Alon, S. and Cui, Y. and Emenari, A. and Goodwin, D. and Karagiannis, E. and Sinha, A. and Wassie, A. T. and Caldas, C. and Bruna, A. and Callari, M. and Greenwood, W. and Lerda, G. and Lubling, Y. and Marti, A. and Rueda, O. and Shea, A. and Harris, O. and Becker, R. and Grimaldi, F. and Harris, S. and Vogl, S. and Joyce, J. A. and Hausser, J. and Watson, S. and Shah, S. and McPherson, A. and Vázquez-García, I. and Tavaré, S. and Dinh, K. and Fisher, E. and Kunes, R. and Walton, N. A. and Sa’d, M. A. and Chornay, N. and Dariush, A. and Solares, E. G. and Gonzalez-Fernandez, C. and Yoldas, A. K. and Millar, N. and Zhuang, X. and Fan, J. and Lee, H. and Duran, L. S. and Xia, C. and Zheng, P. and Marra, M. A. and Hansen, C. and Shah, S. P. (2019) ‘Clonal Decomposition and DNA Replication States Defined by Scaled Single-Cell Genome Sequencing’, cell, 179(5), pp. 1207–1221.e22.

